# A mechanistic model for nuclear migration in hyphae during mitosis

**DOI:** 10.1101/2023.04.12.536534

**Authors:** Subhendu Som, Raja Paul

## Abstract

*S. Cerevisiae* and *C. Albicans*, the two well-known human pathogens, can be found in all three morphologies, i.e., yeast, pseudo-hyphae and true-hyphae. The cylindrical daughter-bud (germ tube) grows very long for true-hyphae, and the cell cycle is delayed compared to the other two morphologies. The place of the nuclear division is specific for true-hyphae determined by the position of the septin ring. However, the septin ring can localize anywhere inside the germ tube, unlike the mother-bud junction in budding yeast. Since the nucleus often migrates a long path in the hyphae, the underlying mechanism must be robust for executing mitosis in a timely manner. We explore the mechanism of nuclear migration through hyphae in light of mechanical interactions between astral microtubules and the cell cortex. We report that proper migration through constricted hyphae requires a large dynein pull applied on the astral microtubules from the hyphal cortex. This is achieved when the microtubules frequently slide along the hyphal cortex so that a large population of dyneins actively participate, pulling on them. Simulation shows timely migration when the dyneins from the mother cortex do not participate in pulling on the microtubules. These findings are robust for long migration and positioning of the nucleus in the germ tube at the septin ring.

## INTRODUCTION

*The opportunistic human pathogens, C. Albicans* and *S. Cerevisiae*, can grow as yeast, pseudo-hyphae and true-hyphae [1–4]. Morphologically these three cell types are different, and also, their cell cycle progressions are not the same [2, 3, 5]. The yeast form is nearly spherical, while pseudo-hyphae is elongated, and true-hyphae resembles a cylindrical shape. In yeast, cell division initiates by forming a ring-like structure of septin on the surface of the mother-bud [6–8]. The septin position is marked as the bud emergence site from where the bud initiates and grows with time [9–11]. The nucleus within the mother-bud migrates to the mother-daughter junction (septin position), and the spindle inside the nucleus becomes parallel to the mother-daughter axis [8, 12, 13]. Then, anaphase is initiated, and the two buds are separated from each other following cytokinesis [14–16]. Pseudo-hyphae cells also follow similar mitotic steps, besides a few exceptions: (a) the bud emerges in a unipolar pattern (from the opposite pole of the birth scar from the previous cell cycle) and upon cytokinesis, the daughter-buds do not separate from the mother; (b) consequently, branched chains of elongated buds are formed [3, 4, 17–21]. Unlike budding yeast, both the mother- and daughter-buds in pseudo-hyphae cells spend a long time in G2 and therefore enter their next cell cycle nearly synchronously [3, 22–24]. The differences between budding yeast and true-hyphae (or simply hyphae) are even more significant. In hyphae, the growth of the daughter-bud is polarized instead of being isotropic [3, 25–30]. As a result, a tubular daughter-bud, called a germ tube, comes out from the mother and over time, it can grow very long [31–35]. This polarized bud growth is directed by a hyphal-specific organelle spitzenkörper which is different from polarisome that regulates isotropic bud growth in yeast and pseudo-hyphae [3, 19, 36–39]. Septin ring forms anywhere inside the germ tube, and the nuclear division takes place at this septin position [3, 40–42]. Like pseudo-hyphae, hyphal buds do not separate after cytokinesis and therefore produce branched chains of hyphal tubes [29, 43–45]. The objective of this study, as described below, aims to explore the mechanistic pathways that are responsible for proper nuclear migration through the germ tube.

Although several genes and motor proteins participate in nuclear migration in hyphae, it is identified that cortical dynein and microtubule primarily regulate the migration [40, 46–49]. Given that mother-bud and hyphae have distinct geometries and possibly different molecular interactions among the primary molecular players, a quantitative understanding of the nuclear migration is demanding. Existing literature hypothesized models based on a) *microtubule (MT) gliding*, b) *dynein ‘pull’ on nucleus*, c) *dynein ‘pull’ on spindle pole bodies (SPBs)* and d) *transport of nucleus as cargo* [40, 46–48]. According to *microtubule gliding model*, cytoplasmic microtubules can lie in an antiparallel configuration in the hyphal tube and slide when dynein is attached between them. Eventually, some microtubules move toward the hyphal tip. The nucleus can bind to the sliding microtubules and gradually migrates along the hyphal tube. The *dynein ‘pull’ on nucleus* allows aggregated dyneins at the hyphal tip to pull on the cytoplasmic microtubules attached to the nucleus. This leads to the migration of the nucleus along the tube. *Dynein ‘pull’ on SPBs* is facilitated by astral microtubules that are nucleated from SPBs and reach the hyphal cortex to interact with dyneins. Since the SPBs are embedded on the nucleus, the dynein pull on the SPBs can move the nucleus along the hyphal tube. *Transport of nucleus as cargo* can be achieved by dyneins directly attached to the nucleus and walking along the cytoplasmic microtubules. If the −ve ends of such microtubules are oriented toward the hyphal tip, nuclear migration can occur. Among all these models, the most significant is the dynein ‘pull’ on SPBs as it is experimentally established for nuclear migration in budding yeast [50–53]. In our earlier work, we considered this model with additional cortical interactions of the astral microtubules to show that the model can unravel many mitotic processes in budding yeast *C. Neoformans* [54, 55]. Using a similar approach, in the present study, we explore the fidelity of nuclear migration in hyphae using numerical simulation. Our data signifies that cortical pushing on the astral microtubules from the mother-bud and pulling from the hyphae are essential for a successful migration.

## RESULTS

### Model and Simulation

Interactions between astral microtubules and the cortex generate three types of forces on the microtubule-tips, viz., pushing force, pulling force and sweeping force [8, 13, 56–63]. Pushing force repeals the tips away from the cortex, whereas pulling force attracts the tips toward the cortex and sweeping force pushes (slides) the tips toward the septin ring along the cortex [8, 13, 56–63]. Pushing force is generated as cortex opposes polymerization of microtubule, pulling force is generated as the cortically anchored dyneins bind the microtubule and walk along the filament toward -ve and sweeping force is generated as cortical bim1-kar9-myo2 complex is formed on the microtubule-tip sliding toward the septin ring [8, 13, 56–63]. Among the three forces, sweeping force is mother-bud specific [8, 13, 60–63] and the other two forces are active in both the mother and daughter cortices [56, 58, 59]. In this study, cortex is considered ∼ 0.2 *µm* wide from the cell-membrane [7, 8, 55].

Mother-bud, nucleus and the SPBs are considered as spherical objects [54, 55] of radii *r*_*M*_, *r*_*nuc*_ and *r*_*spb*_, respectively. Daughter-bud is a cylinder of radius 1.5 *µ*m [27, 29] and its polarized elongation is very much (here 9 *µ*m) [3, 5] (Fig. 1). Initially, the nucleus is randomly placed in the mother-bud and the SPBs are embedded on the nuclear envelope [8, 54]. At this stage, the SPBs are kept very close to each other. Dynein motors are distributed in the cortical region. The septin ring is placed halfway through the daughter-bud. Sister kinetochores (KTs) are taken as spheres [8, 54, 55] of uniform radius *r*_*kt*_ and distributed in the nucleus. Microtubules that connect kinetochores with the SPBs are known as kinetochore microtubules (kMTs). Kinetochores form amphitelic attachments with the two SPBs (Fig. 1). For simplicity, we do not consider monotelic, syntelic and merotelic attachments of the sister kinetochores. Cohesin binds the sister kinetochores until the beginning of anaphase and behaves like a spring [8, 54, 55]. Due to polymerization and depolymerization of the kMTs while interacting with the kinetochores (Fig. 1), attractive and repulsive forces are generated on the microtubule-tips [8, 54, 55, 64, 65]. These forces contribute to the movement of the two SPBs over the nuclear envelope [8, 54, 55]. The kinesin5 motors bind the anti-parallel microtubules (interpolar microtubules) of the two SPBs (Fig. 1) and produce repulsive force between the SPBs [8, 54, 55, 66]. This is the primary force within the nucleus that facilitates SPB separation. The kinetochore microtubules and the interpolar microtubules are collectively called nuclear microtubules (nMTs). The structure of the metaphase spindle depends on the nuclear microtubules. Besides nuclear microtubules, the two SPBs nucleate another set of microtubules which are found in the cytoplasm and interact with the cell cortex [8, 12, 67, 68]. These microtubules are known as astral microtubules (aMTs). Due to cortical interactions of the astral microtubules, three types of forces are generated on the microtubule-tips and they are finally transmitted to the center of mass of the nucleus and therefore responsible for the nuclear movement in the cell. These three forces are cortical pushing, pulling and sweeping forces (see above). The mechanisms by which the forces are produced are described below.

**FIG. 1.**
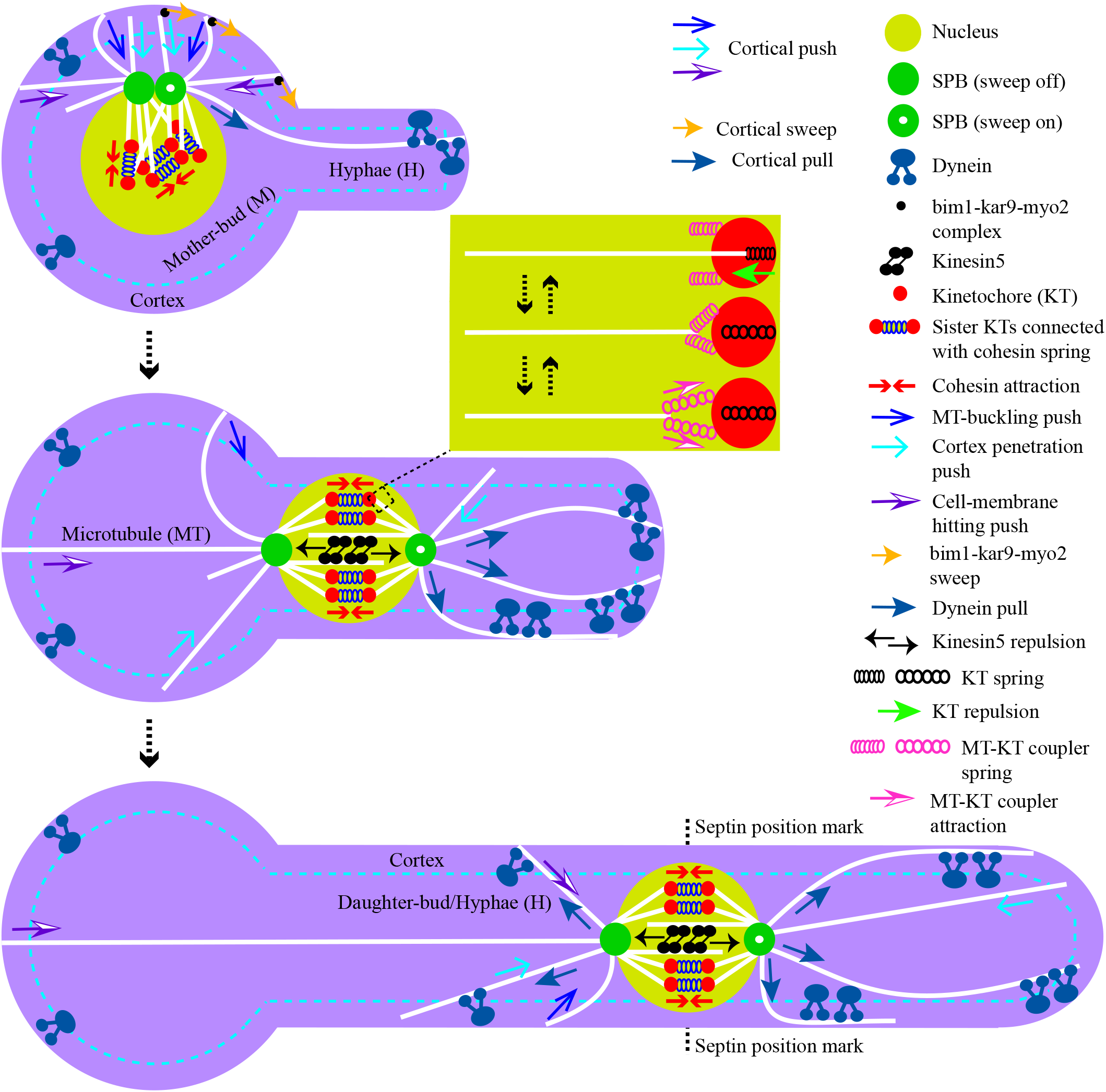
Model schematic used to simulate nuclear movement in the hyphae cells. Various intracellular components and forces considered in the model are shown. Daughter-bud (Hyphal tube) grows with time and the nucleus reaches near the septin region driven by forces arising from the interactions between the astral microtubules and cortex. The mitotic spindle assembles within the nucleus and orients parallel to the hyphal tube.

### Cortical pushing forces

When a polymerizing microtubule penetrates the spring-like cortex, spring pushing force 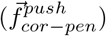 is applied on the tip [56, 57]. If the cortical penetration length of the microtubule is *l*_*cor*−*pen*_ and the cortical stiffness is *κ*_*cor*−*pen*_, the force would be 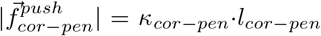. After entering into the cortex, if the microtubule continues polymerizing and bumps into the cell membrane, an instantaneous pushing force (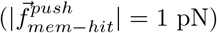) is applied on the tip [8, 54, 55, 69]. Reaching the cell-membrane, the microtubule can either buckle [54, 70, 71] or undergo depolymerization [54, 72, 73] with equal probabilities. For a buckled microtubule, we consider first-order Euler’s buckling force i.e. 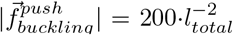 [54, 69, 74, 75]. Here *l*_*total*_ is the microtubule length. Notably, the buckling force is directed along the imaginary line connecting the microtubule-tip and SPB. Since all these three forces can be generated in both mother- (M) and daughter- (D) buds, force terms and the parameters are represented with appropriate sub/super-scripts; e.g., 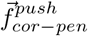 would be 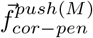 and 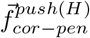, if 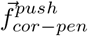 is generated in mother- and daughter-buds, respectively.

### Cortical pulling force

Cortically anchored dynein motors bind the +ve end of the microtubule and in a bid to walk toward the −ve end of the filament they keep pulling on the microtubule [58, 59]. The force generated is of the Form 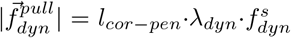, where *l*_*cor−pen*_ is cortical penetration length of microtubule, *λ*_*dyn*_ is cortical dynein density and 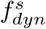 is the single dynein force [8, 54, 55, 69].

### Cortical sweeping force

Cortical bim1, kar9 and myo2 proteins form a complex (bkm complex) that sweeps microtubule-tip toward the septin ring along the cortex [8, 13, 60–63]. If *l*_*cor*−*pen*_ is the cortical penetration length of the microtubule, *λ*_*bkm*_ is cortical bkm density and 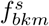 is single bkm force, the sweep force on microtubule is 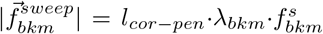. Note that sweep force is mother-bud specific and also specific for the microtubules originating from one of the SPBs (see the schematic of Fig. 1) [8, 13, 60–63]. Therefore when the microtubules from the designated SPB leave the mother cortex, the sweep force vanishes.

### Movement of the nucleus

Since the nucleus travels through viscous cytoplasm, we use Stokes’ law to update nuclear position with time [8, 54, 55, 69, 75, 76]. If 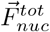 is the net instantaneous force acting on the nucleus causing a velocity 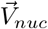, then 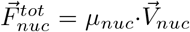. Here *µ*_*nuc*_ is the coefficient of effective viscous drag applied on the nucleus and 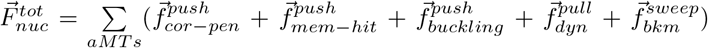. Since 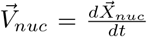, once we know instantaneous 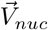, nuclear position 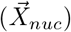 can be updated time. In the simulation, the time step (∆t) is 0.01 s.

### Kinetochore forces

When a kinetochore microtubule polymerizes against the kinetochore (see Fig. 1), pushing force applied on the microtubule-tip [8, 54, 55]. If *κ*_*kt*_ is the stiffness and *∆l*_*kt*_ is the compressed spring length of kinetochore then 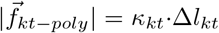. Since 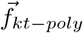 is a load force on the microtubule, it will undergo depolymerization and leave the kinetochore (Fig. 1). While leaving the kinetochore, coupling proteins residing on the kinetochore may attach with the depolymerizing microtubule and pull it toward the kinetochore [8, 54, 55]. If *κ*_*coupler*_ is the stiffness and *∆l*_*coupler*_ is the stretched length of the coupling proteins pulling force on the microtubule-tip would be 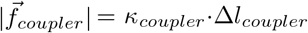.

### Kinesin5 force

In the simulation, we do not explicitly consider the kinesin5 binding to the interpolar microtubules, rather an equivalent force expression is used to generate repulsive force between the two SPBs [77, 78]. If *d*_*spb*−*spb*_ and *l*_*av*_ are SPB-SPB distance and average interpolar microtubule length, respectively, sliding (repealing) force acting on the microtubule-tips is 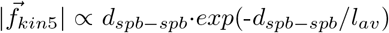.

### Equation of motion of SPB

SPBs move over the nuclear envelope and therefore the motion is overdamped and we use Stokes’ law to update SPB position with time [8, 54, 55]. If 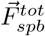 is the instantaneous net force acting on the SPB, 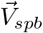 is the instantaneous velocity of SPB and *µ*_*spb*_ is the effective viscous drag on SPB then 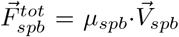. Here 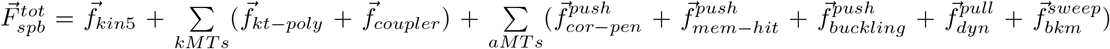 is provided in the Table I. Since 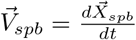 with time, once we know 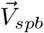, instantaneous SPB position 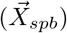 can be updated with time.

**TABLE 1.** Model Parameters.

| Abbreviations | Meanings | standard values | References |
| --- | --- | --- | --- |
| $r_M$ | Mother-bud radius | $3 \mu m$ | [8, 54, 55] |
| $r_{nuc}$ | Nuclear radius | $1 \mu m$ | [8, 54, 55] |
| $r_{spb}$ | SPB radius | $0.125 \mu m$ | [8, 54, 55, 97, 98] |
| $r_{kt}$ | Kinetochore radius | $0.05 \mu m$ | [8, 99] |
| $v_{g0}^{aMTs}$ | Polymerization speed of astral microtubules | $6.4 \mu m \cdot min^{-1}$ | [8, 100, 101] |
| $f_{c0}^{aMTs}$ | Catastrophe rate of astral microtubules | $0.34 min^{-1}$ | [8, 100, 101] |
| $v_s^{aMTs}$ | Depolymerization speed of astral microtubules | $26.6 \mu m \cdot min^{-1}$ | [8, 100, 101] |
| $f_r^{aMTs}$ | Rescue rate of astral microtubules | $0.02 min^{-1}$ | [8, 54, 55, 100, 101] |
| $f_c^{stall(aMTs)}$ | Catastrophe rate of stalled astral microtubules | $0.04 s^{-1}$ | [54, 55, 76] |
| $f_{stall}^{aMTs}$ | Stall force of astral microtubules | $1.7 pN$ | [8, 54, 55, 81] |
| $v_{g0}^{nMTs}$ | Polymerization speed of nuclear microtubules | $6.4 \mu m \cdot min^{-1}$ | [102] |
| $f_{c0}^{nMTs}$ | Catastrophe rate of nuclear microtubules | $2.14 min^{-1}$ | [102] |
| $v_s^{nMTs}$ | Depolymerization speed of nuclear microtubules | $18.6 \mu m \cdot min^{-1}$ | [102, 103] |
| $f_r^{nMTs}$ | Rescue rate of nuclear microtubules | $0.04 min^{-1}$ | [102] |
| $l_{av}$ | Average interpolar microtubule length | $\sim 3 \mu m$ | [75] |
| $\kappa_{cor-pen}$ | Rigidity of cortex | $5 pN \cdot \mu m^{-1}$ | [8, 54, 55] |
| $\lambda_{dyn}$ | Cortical dynein density | $6 \mu m^{-1}$ | [8, 54, 55] |
| $f_{dyn}^s$ | Single dynein force | $1 pN$ | [8, 54, 55] |
| $\lambda_{bkm}$ | Cortical BKM density | $6 \mu m^{-1} = \lambda_{dyn}$ | [54, 55] |
| $f_{bkm}^s$ | Single BKM force | $1 pN$ | [55] |
| $\kappa_{kt}$ | Stiffness of kinetochore | $5 pN \cdot \mu m^{-1}$ | [8, 54, 55, 65, 104] |
| $\kappa_{coupler}$ | Rigidity of kinetochore-microtubule couplers | $10 pN \cdot \mu m^{-1}$ | [8, 54, 55, 65, 104] |
| $\mu_{nuc}$ | Viscous cytoplasmic drag on nucleus | $\sim 100 pN \cdot s \cdot \mu m^{-1}$ | [8, 54, 55] |
| $\mu_{spb}$ | Viscous drag of nuclear envelope on SPBs | $\sim 25 pN \cdot s \cdot \mu m^{-1}$ | [8, 54, 55] |
| $\mu_{kt}$ | Viscous drag of nucleoplasm on kinetochores | $\sim 10 pN \cdot s \cdot \mu m^{-1}$ | [8, 54, 55] |
| $N_d$ | Maximum permissible number of dyneins on astral microtubules | $12 \mu m^{-1}$ | [55, 84, 85] |
| $K_{on}$ | Dynein attachment rate to astral microtubules | $1.6 s^{-1}$ | [55, 84, 85] |
| $k_{u0}$ | Dynein detachment rate from astral microtubules | $0.27 s^{-1}$ | [55, 84, 85] |
| $F_d$ | Dynein detachment force for astral microtubules | $0.67 pN$ | [55, 85] |

### Cohesin force

Sister kinetochores are joined by spring-like cohesin having rest length 0.1 *µm* and spring constant 0.1 pN · µ*m*^−1^ [8, 54, 55, 79]. Due to the movement of the kinetochores during spindle formation, if cohesin length changes 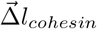, a force 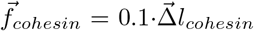 is applied on the kinetochores.

### Equation of motion of kinetochore

Kinetochores move through viscous nucleoplasm, so the motion is over-damped. Like the nucleus and SPBs, Stokes’ law is followed to update kinetochore position with time [8, 54, 55]. If 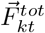 is the instantaneous net force acting on the kinetochore, 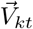 is the instantaneous velocity of kinetochore and *µ*_*kt*_ is the effective viscous drag on kinetochore then 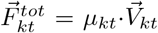. Here 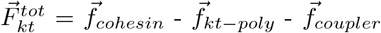 and *µ*_*kt*_ is provided in the Table I. Integrating 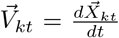, instantaneous kinetochore position 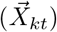 can be updated.

### Steric forces

Steric repulsion is present between overlapping i) nucleus and cell-membrane, ii) kinetochore and nuclear envelope and iii) non-sister kinetochores.

The forces (except 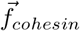 and steric) described above are microtubule-dependent. Microtubules are dynamic polymers and their dynamic instability is reproduced in simulation using four parameters [8, 54, 55, 69, 75, 76]: polymerization speed *v*_*g*_, the rate of transition from polymerization state to depolymerization (catastrophe frequency) *f*_*c*_, depolymerization speed *v*_*s*_ and rate of recovery from depolymerization state to polymerization (rescue frequency*) f*_*r*_ [8, 54, 55, 69, 75, 76]. Note that aMTs and nMTs have different sets of dynamic instability parameters (*v*_*g*_, *f*_*c*_, *v*_*s*_ and *f*_*r*_) (Table I). If a microtubule-tip is pushed, its *v*_*g*_ and *f*_*c*_ are modified as: *v*_*g*_ *= v*_*g0*_ exp(−*f*_*load*_*/f*_*stall*_) and 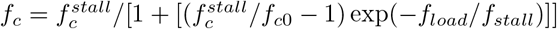, where, *f*_*load*_ is the pushing load, *f*_*stall*_ is the load that stalls the growth of a single microtubule, *v*_*g0*_ is the unconstrained polymerization speed under zero load, 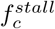 is the rate of catastrophe of a stalled microtubule *and f*_*c0*_ is the catastrophe rate of a free microtubule [8, 54, 55, 69, 76, 81]. In our study, we consider four aMTs and twenty nMTs per SPB [8, 55, 80, 82, 83]. Among twenty, sixteen are the kMTs and four are the interpolar microtubules.

In this study, nuclear migration time is defined as the average time taken by the nucleus to reach the septin ring. Each simulation runs for a long duration (*∼* 300*min*) and a nucleus reaching the septin ring within this time is considered a success, otherwise, the migration is recorded as a failure. In the figures (Fig. 2 - Fig. 5), scale-break is given (e.g. >300 min) to denote the failed migrations.

**FIG. 2.**
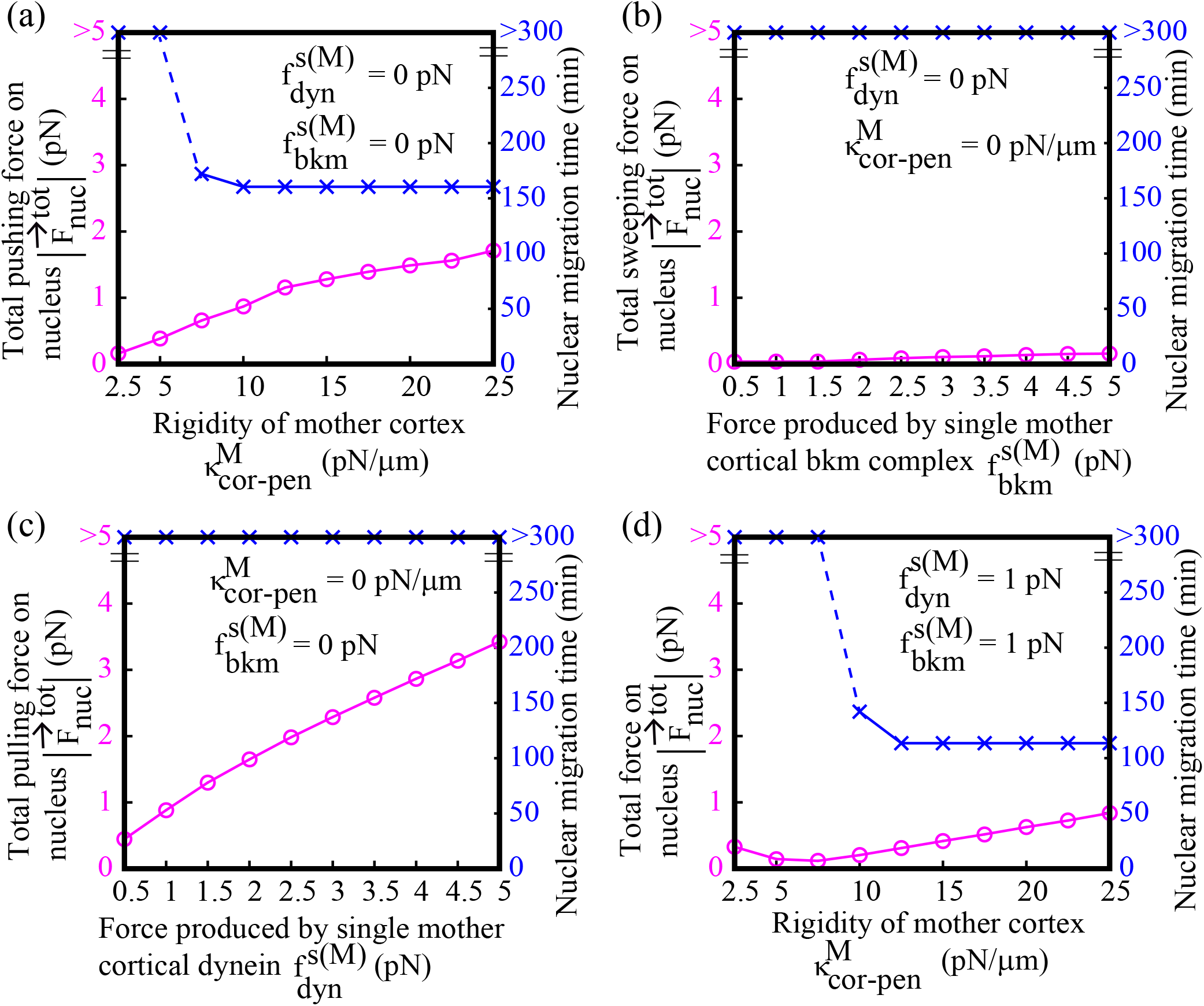
Cortical pushing forces originated in mother-bud promote nuclear migration. In the absence of all forces produced in daughter-bud (a-d), cortical pushing forces from the mother-bud facilitates nuclear migration to the hyphal tube within finite time (a and d). (a) Among the three cortical pushing forces (i.e. 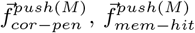 and 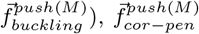 is considered to study the effect of cortical pushing on the nuclear migration. With the increase of 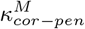, the total pushing force from the mother cortex on the nucleus increases reducing migration time. (b) If the cortical sweep (bkm force) works alone in the mother-bud, the required force for nuclear migration is not achieved resulting in unsuccessful migration. (c) When cortical pull (dynein force) independently applies on the astral microtubules from both the SPBs, the nucleus moves toward the mother cortex opposing nuclear migration toward the hyphal tube. (d) For higher values of 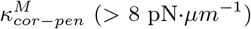 (> 8 pN*·µm*^−1^), cortical pushing overcomes cortical pulling promoting successful migration.

### Stochastic dynein density in cortex

Among the three cortical forces, cortical pulling 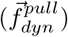 depends on the dynein density (*λ*_*dyn*_) in cortex. We first study nuclear migration under constant dynein density (6 *µm*^−1^). However, practically, local dynein density is stochastic and may depend on conditions such as the maximum permissible number of dyneins on microtubule *N*_*d*_, dynein attachment rate to microtubule *K*_*on*_, dynein detachment rate from microtubule *k*_*u0*_ and dynein detachment force for microtubule *F*_*d*_) [55, 84, 85]. The implementation of stochastic dynein density is described below.

If *n(t)* denotes the instantaneous number of dyneins attached with a microtubule (*n* <= *N*_*d*_), the rate at which any unbound dynein binds to the microtubule is *k*_*on*_*(n)* = *(N*_*d*_ − *n) · K*_*on*_ [55, 84, 85]. Following Bell’s (or Kramers’) theory, we assume that the instantaneous rate of unbinding one of the n dyneins from the microtubule is approximately 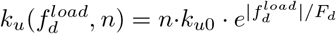. Here 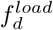 is the load force on the dynein. For simplicity, we assume the total pushing force (load) generated on the microtubule-tip is equally shared by the *n* dyneins [55, 84]. At each time step of the simulation, we use the above two expressions to determine the local dynamic dynein density in the cortex.

In order to test the fidelity of the cortical forces toward the nuclear migration, we proceed with various combinations of the forces as follows. Contribution of forces arising from the mother-bud (Fig. 2) and hyphal tube (Fig. 3, top panel) are checked independently, followed by a combined effect (Fig. 3, bottom panel). We explored the effect of a constant cortical dynein density (Fig. 2 and Fig. 3) and stochastic dynein density (Fig. 4) on nuclear migration. Since cortical dynein pulling from the mother-bud opposes nuclear migration (Fig. 2c), we completely suppress the activity of mother cortical dyneins and check nuclear movement (Fig. 5). To test the robustness of our model, we vary the position of the septin ring and follow the nuclear movement (Fig. 5f). The outcome identifies the pathways to a finite nuclear migration time and right spindle orientation for different positions of the septin ring in the hyphal tube.

**FIG. 3.**
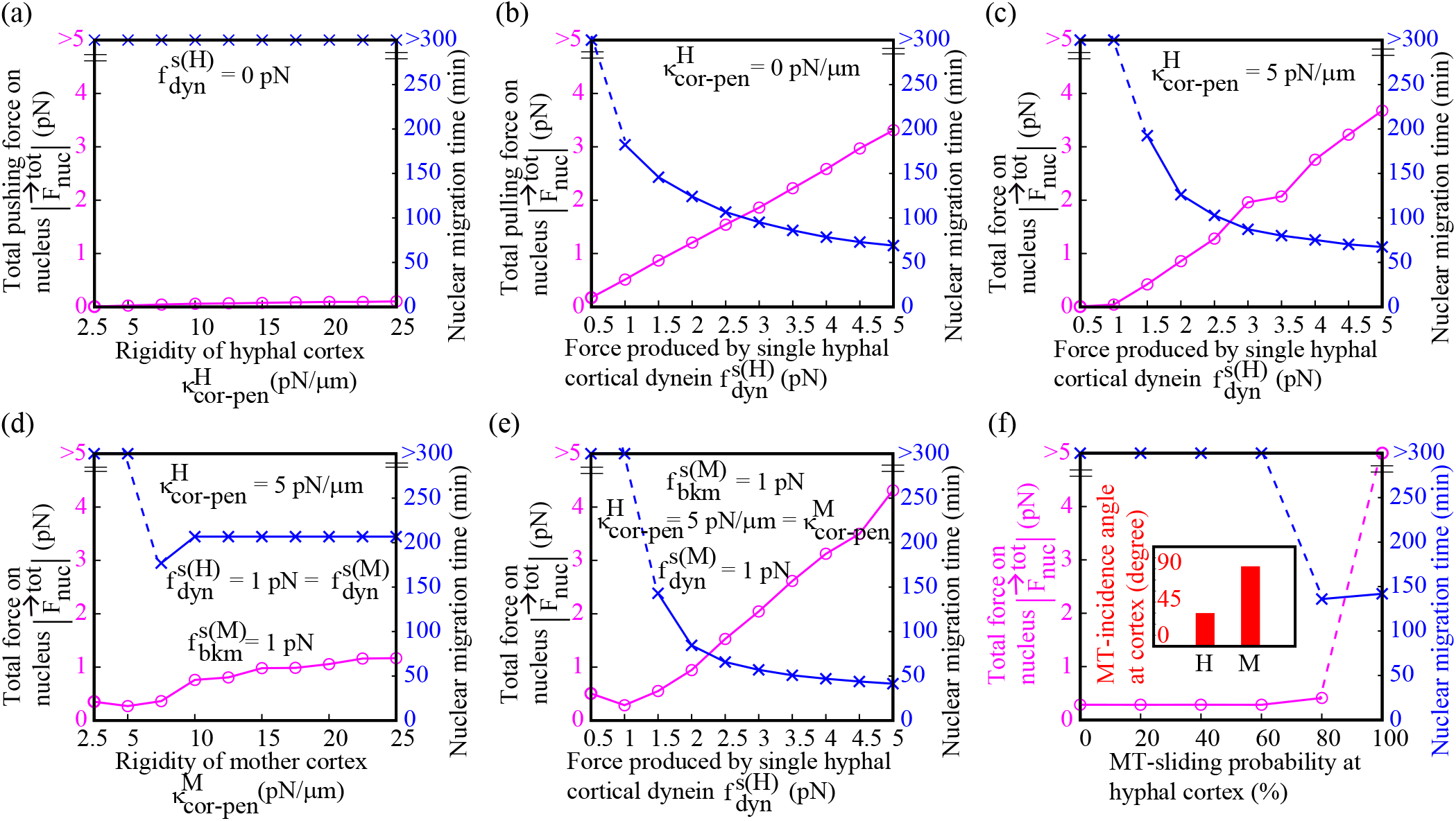
Strong cortical pull from the hyphal tube is necessary for effective nuclear migration. In the absence of all forces produced in mother-bud (a-c), the cortical pulling force from daughter-bud helps the nucleus to reach the septin ring within finite time (b and c). (a) Among the three cortical pushing forces (i.e. 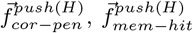 and 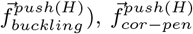 is considered to study the effect of cortical pushing on the nuclear migration. With the increase of 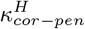, the total pushing force from the hyphal cortex on the nucleus increases opposing nuclear migration toward the hyphal tube. (b) If the cortical pull (dynein force) works alone in the daughter-bud, the required force for nuclear migration is achieved by the nucleus and therefore migration occurs within a finite time. (c) The presence of cortical pushing forces suppresses cortical pulling force for small values of dynein force (<= 1 pN) extending nuclear migration time. Increased dynein force enables cortical pull to dominate over cortical push resulting in finite migration time. (d and e) Cortical pushing from mother-bud and cortical pulling from the daughter-bud (similar to Fig. 2a and Fig. 3b, respectively) can migrate the nucleus when all other forces from the mother- and daughter-buds are active. With the increase of 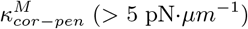 (> 5 pN*·µm*^−1^) and single dynein force (> 1 pN), a finite migration time is observed. (f) In the combined force-field scenarios (d and e), if the microtubules of right SPB (as in Fig. 1) frequently slide along the hyphal membrane, several dyneins can be attached to them and generate strong cortical pull on the nucleus. Thus, finite nuclear migration time is observed at smaller values (standard) of 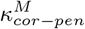 (5 pN·*µm*^−1^) and single dynein force (1 pN).

**FIG. 4.**
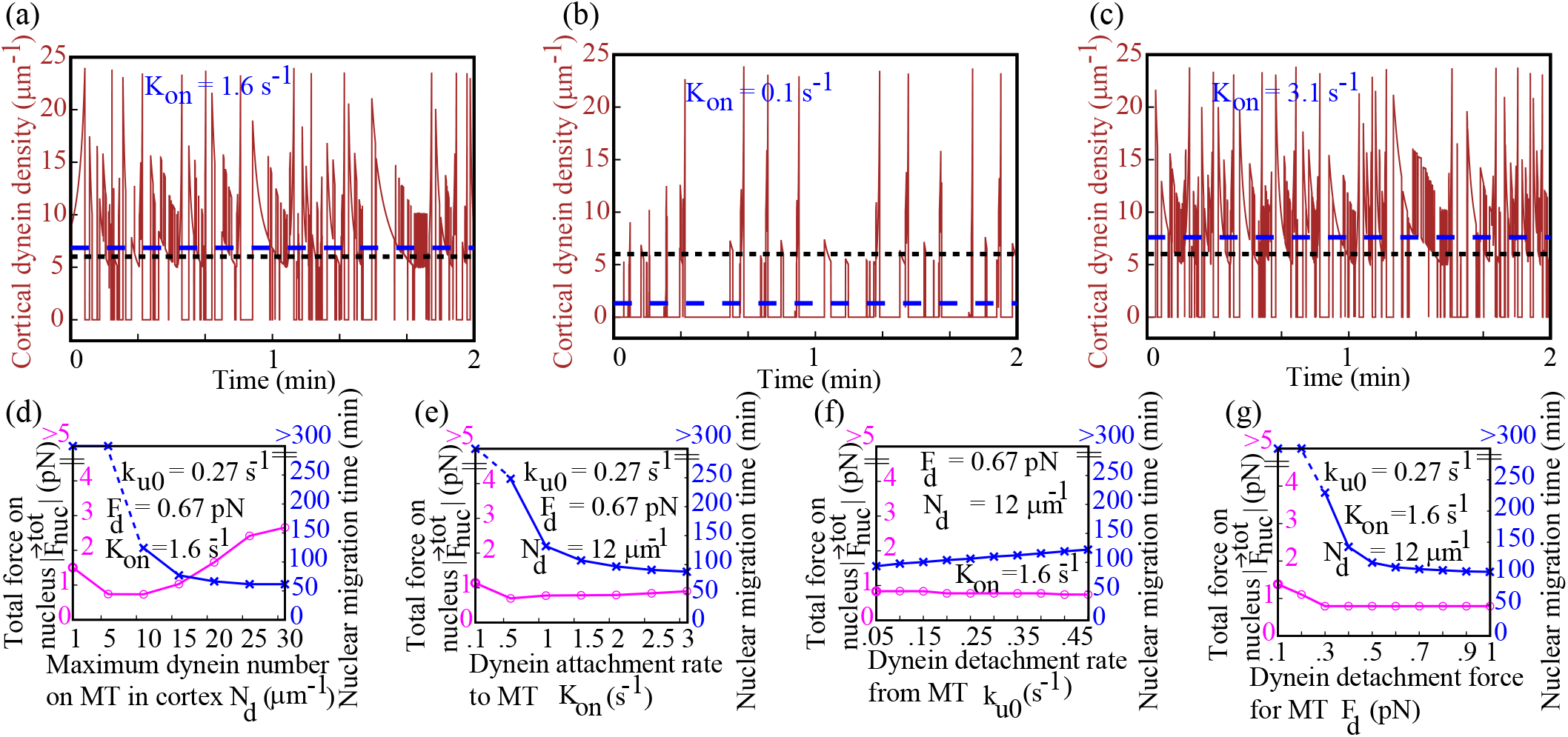
Stochastic dynein binding to microtubule confirms significant cortical pull from the hyphal tube. The requirement of cortical pulling from the hyphal tube for successful nuclear migration is re-investigated by substituting constant dynein density with stochastic dynein density in the cortex. (a-c) Temporal change of cortical dynein density is shown when dynein attachment to microtubule is stochastic (brown line) instead of deterministic (blue dashed line). The average of stochastic dynein density is indicated with a black dotted line. *K*_*on*_ is the rate at which dynein binds microtubule. (d-g) Parameters that control stochastic dynein density (*N*_*d*_, *K*_*on*_, *k*_*u0*_, *F*_*d*_) are sequentially varied, and nuclear migration time is measured. Increase of *K*_*on*_, *N*_*d*_ and *F*_*d*_ and decrease of *k*_*u0*_ lead to early migration. The data presented here (d-g) validate it. For lower values of *K*_*on*_, N_*d*_ and F_*d*_, cortical pulling from the hyphal tube becomes smaller and nuclear migration is inefficient.

**FIG. 5.**
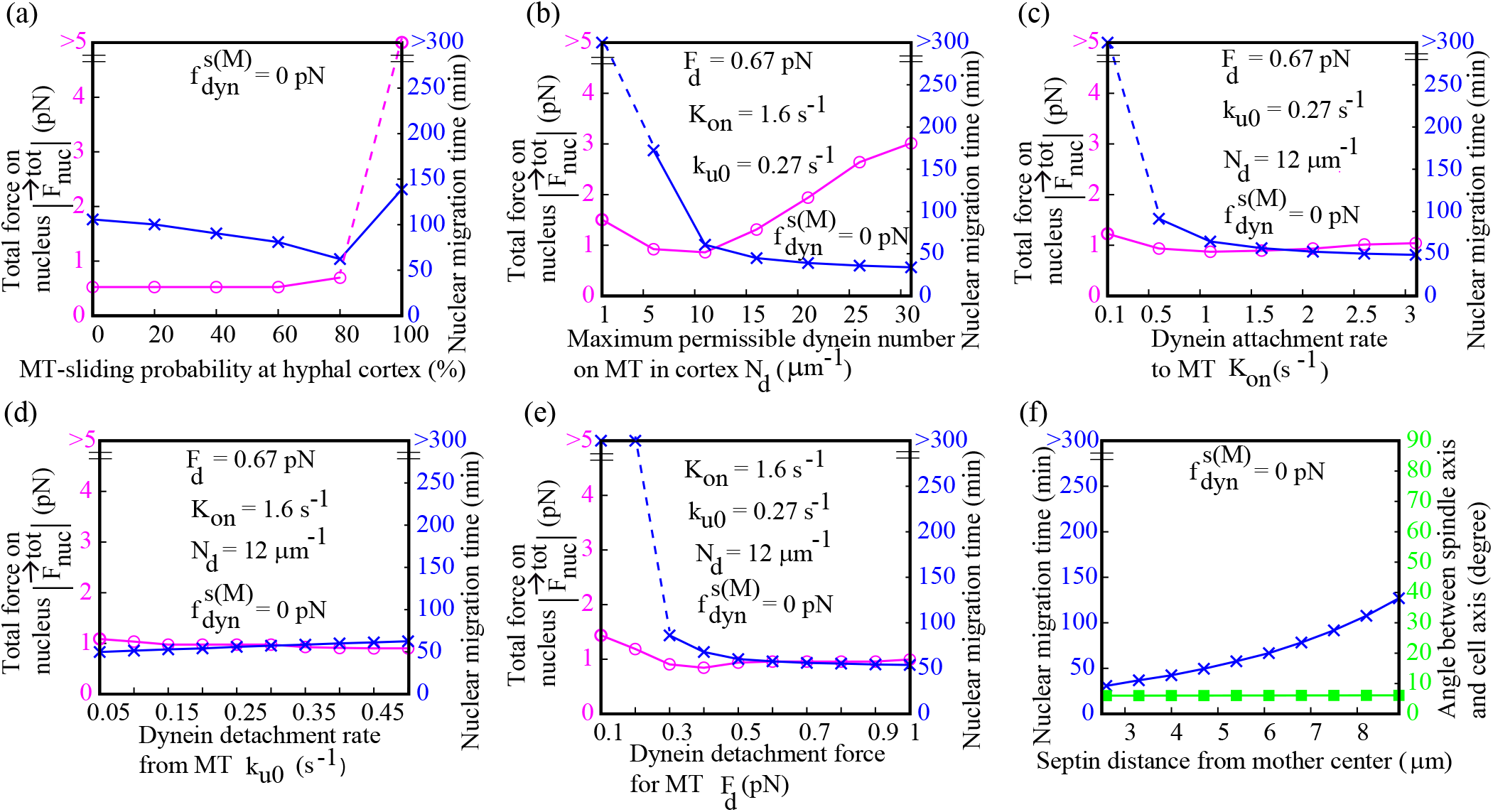
Absence of cortical pulling from mother-bud strongly favors nuclear migration. (a) Similar to Fig. 3f, however, cortical pulling from the mother-bud is absent. The comparative study indicates that cortical pulling from mother-bud strongly opposes nuclear migration. (b-e) Revisiting Fig. 4d to Fig. 4g in the absence of cortical pulling from mother-bud. Migration time is highly reduced if cortical pulling from mother-bud is off. (f) The position of the septin ring is varied in the hyphal tube to study nuclear migration time. Except for cortical pulling from mother-bud, all the forces from both buds apply on the nucleus. It is clear that for any position of the septin ring, finite migration time is achieved and the spindle axis aligns with the cell axis.

### Cortical pushing force from mother-bud can be significant for nuclear movement toward hyphal tube

A polymerizing microtubule in contact with the mother cortex, experiences three types of pushing force i.e. 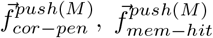 and 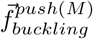 (see Model; Fig. 1). Here we quantify the effect of these pushing forces on nuclear migration by omitting forces generated in daughter-bud. We increase rigidity of the cortex 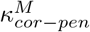 gradually and measure net force on the nucleus 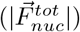. Fig. 2a shows that with the increase of 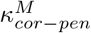, net force on the nucleus 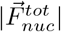 increases monotonically while the nuclear migration time decreases abruptly and then saturates. During variation of 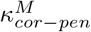, all other pushing forces at the mother cortex, such as instantaneous pushing force from the membrane and force due to microtubule buckling are kept constant at standard values. We identify that most of the microtubules from the left SPB (schematic of Fig. 1) engage with the mother cortex and generate the pushing force, whereas some of the microtubules from the right SPB interact with the daughter cortex. Therefore, the net cortical pushing on the left SPB is greater than the right SPB. As a result, the nucleus moves rightward in the daughter-bud.

Next, we focus on the independent contribution of the cortical sweep 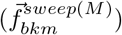 on the migration (Fig. 2b). We set all other forces to zero and vary the force due to the bkm complex. bkm force (i.e., the bias force) is mother-bud specific and also specific to microtubules from one particular SPB (the right SPB as per Fig. 1). Therefore when the microtubules from the right SPB do not encounter the mother cortex, bkm force dies out. Consequently, the net bias force on the nucleus becomes insignificant and nuclear migration is not achieved.

Further, we discuss the independent contribution of the cortical pulling from the mother-bud 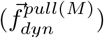 on the nuclear migration in Fig. 2c. Similar to the cortical pushing scenario, the net cortical pulling on the left SPB is greater than the right SPB due to more microtubules from the left SPB interacting with the mother cortex. As a result, the nucleus tends to move toward the mother cortex, i.e., nuclear migration is opposed by the pulling from the mother cortex.

Since only the cortical pushing force promotes nuclear migration, we re-investigate the contribution of the pushing force on migration when cortical sweeping and pulling are present in the mother-bud (Fig. 2d). The data show that the pushing force is capable of producing a finite migration time together with other forces. When the cortical rigidity parameter is 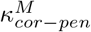 is small, cortical pulling dominates over cortical pushing and sweeping forces and therefore migration is impaired. At larger 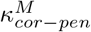, the pushing force aided by the sweeping force exceeds the resistance of the pulling from the mother cortex and achieves timely nuclear migration.

### Cortical pulling force from hyphal tube favors nuclear migration

Before reporting the results due to pulling force we discuss the effect of cortical pushing forces 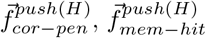 and 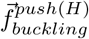 from the hyphal tube. Microtubules penetrating the cortex translate the pushing force 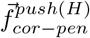 on the nucleus that depends on the rigidity parameter 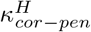 as shown in Fig. 3a. In the simulation, pushing 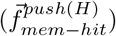 and buckling 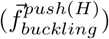 forces are generated on the microtubule-tips using the standard parameters and rest of the forces from the hyphal tube (e.g., dynein pull 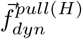 are ignored. To measure the independent effect of the hyphal tube on nuclear migration, all the mother-bud forces are turned off (Fig. 3a - Fig. 3c). The schematic in Fig. 1 shows, while the microtubules from the right SPB easily reach the hyphal cortex, microtubules from the left SPB cannot reach there. Consequently, the net pushing force on the right SPB is transmitted to the nucleus shifting leftward i.e. migration is not favored in this scenario.

To check the independent contribution of cortical pulling force 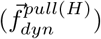 on nuclear migration, we ignore all the cortical pushing forces and measure net pull on the nucleus by varying the single dynein force (Fig. 3b). Net pulling on the right SPB is transmitted to the nucleus and it migrates rightward. After entering the hyphal tube, microtubules from both the SPBs interact with the hyphal cortex and apply an opposite pull on the nucleus. While the cortical interactions of the microtubules from the right SPB remain unaltered, some of the microtubules from the left SPB reach the mother-bud and do not interact with the hyphal cortex (Fig. 1). Hence net cortical pull on the nucleus remains rightward and migration continues in the hyphal tube. In Fig. 3c, we check the efficiency of the cortical pulling force corresponding to the finite migration time when the cortical pushing forces are turned on. It is observed that the contribution of cortical pulling on migration is almost the same without the cortical pushing force as shown in Fig. 3b. For smaller values of dynein force (<= 1 pN), cortical pushing dominates cortical pulling and therefore migration is perturbed.

Results shown in Fig. 2 and the top panel of Fig. 3, suggest that cortical pushing from the mother-bud and cortical pulling from the daughter-bud are redundant forces that can independently facilitate the nuclear migration. Considering a realistic situation, we now consider all the forces from mother- and daughter-buds and discuss the nuclear migration features in Fig. 3d and Fig. 3e. It is observed that if the rigidity of the mother cortex 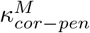 is larger than 5 pN*·µm*^−1^, cortical pushing force from mother-bud is still capable of sending the nucleus to the septin ring. Alternatively, if the single dynein force at the daughter cortex increased beyond 1 pN, cortical pulling from the hyphae can lead to significantly reduced nuclear migration time. Therefore, cortical pushing from mother-bud and cortical pulling from the hyphal tube together can drive the nucleus to the hyphal tube. Although finite nuclear migration time is achieved by combining the forces shown in Fig. 3d and Fig. 3e, 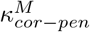 and single dynein force are larger than their respective standard values (see Table I).

Our goal is to explore the potential mechanism that functions with the standard model parameters and gives finite migration time. Therefore we review the mechanism through which a polymerizing microtubule interacts with the hyphal cortex. Previously, a polymerizing microtubule would either buckle at the hyphal membrane or depolymerizes with equal probabilities. Nevertheless, polymerizing microtubules largely slide along the hyphal membrane due to relatively narrow cylindrical geometry where microtubules often hit the cortex at a grazing angle [88–90]. In our further study in Fig. 3f, we consider the sliding probability in addition to the buckling and depolymerizing probabilities. Note that the sum of these three probabilities is unity and during variation of sliding probability, the two other two probabilities are kept the same. From the data shown in Fig. 3f, it is clear that if the microtubule-sliding probability is large at the hyphal cortex (> 60%), finite nuclear migration time can be achieved for the standard values of all the simulation parameters. On a sliding microtubule in the hyphal cortex, several dyneins can be attached and they generate a strong cortical pull which helps the nucleus to migrate to the septin ring for the low values of 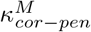 and single dynein force. Sliding microtubule-based large cortical pulling from daughter-bud is experimentally identified for successful nuclear migration in *S. Cerevisiae* yeast [50, 51]. Also, the average angle of incidence of microtubule measured in our simulation is small for the hyphal cortex and large (*∼* 90^0^) for the mother cortex (inset; Fig. 3f).

Thus microtubules slide more frequently in the hyphal cortex than buckling or depolymerizing.

### Stochastic dynein attachment to microtubule validates the positive contribution of cortical pulling from hyphae for nuclear migration

Results shown in Fig. 3 suggest that strong cortical pull on the nucleus from the hyphal tube is necessary for finite migration time. The origin of this force is due to cortical dynein which is, so far, assumed to maintain a constant average density (6 *µm*^−1^). In other words, if the cortical segment of a microtubule is 1 *µm*, six cortical dyneins would always attach to the microtubule-tip. A similar assumption made in earlier studies successfully captured relevant cellular processes [8, 54, 55, 69]. However motor attachment to microtubule is not deterministic, rather it is stochastic and depends on four conditions: i) maximum permissible number of dyneins that can bind the cortical segment of the microtubule (*N*_*d*_), ii) dynein attachment rate to microtubule (*K*_*on*_), iii) dynein detachment rate from microtubule (*k*_*u0*_) and iv) dynein detachment force from microtubule (*F*_*d*_) [55, 84, 85]. The stochastic dynein interaction with microtubules is described earlier in the Model and Simulation section. Here, we explore how the above-mentioned parameters impact the outcome obtained with fixed dynein density (Fig. 3f). We simulate the system with standard parameter values and microtubules sliding along the hyphal cortex with 80% probability (Fig. 4a - Fig. 4g). The four dynein parameters are varied systematically (Fig. 4d - Fig. 4g), i.e. when one parameter is varied, the other three parameters are kept at standard values (see Table I). To elucidate the difference between constant dynein density and stochastic dynein density, we first show the fluctuation of the cortical dynein density with time (Fig. 4a - Fig. 4c). The dotted lines (black) indicate constant dynein density (6 *µm*^−1^), whereas the dashed lines (blue) show the average of the stochastic dynein density. Notice that the average value increases with the increase of dynein attachment rate to the microtubule (*K*_*on*_). Consequently, we see that the nuclear migration time becomes finite and then decreases with the increase of *N*_*d*_, *K*_*on*_ and *F*_*d*_ (Fig. 4d - Fig. 4g). Increase of *N*_*d*_, *K*_*on*_, *F*_*d*_ corresponds to a robust interaction of dyneins with microtubules. This effectively indicates a strong cortical pull from the hyphal cortex on the nucleus leading to a finite migration time. For smaller values of *N*_*d*_ (<= 6 *µm*^−1^), *K*_*on*_ (<= 0.1 s^−1^) and F_d_ (<= 0.2 pN), migration promoting forces (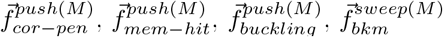 and 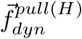) are dominated by the migration suppressing forces (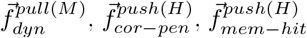 and 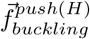) and thus the nucleus mostly remain within the mother-bud. Fig. 4f emphasizes the monotonous increase of the migration time with the increase of dynein detachment rate *k*_*u0*_ implying weaker cortical pulling on the nucleus from the hyphal tube.

### Cortical pulling from mother-bud might be suppressed during nuclear movement toward the hyphal tube

As mentioned in the previous section, forces produced within the mother- and daughter-buds can promote or suppress nuclear migration. Naturally, if the migration suppressing forces are removed, successful nuclear migration would be prevalent and migration time would be less. Interestingly for *S. Cerevisiae* yeast, several articles suggested that the functions of mother cortical dyneins generating pulling forces on the microtubule-tips 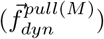, remain off till the nucleus reaches the septin ring [67, 68, 91–93]. In our simulation, we apply this mechanism and show the results in Fig. 5. Comparing Fig. 5a and Fig. 3f, we find that the absence of cortical pulling from the mother-bud sharply decreases migration time. Similar to Fig. 4(d-g), we establish the decreasing migration time by substituting constant dynein density with stochastic dynein density in the cortex (Fig. 5b - Fig. 5e). Comparing Fig. 5(b-e) with the corresponding Fig. 4(d-g), we further see that migration time is lower when cortical pulling from the mother-bud is absent. From these results, we suggest that possibly for hyphae the cortical pulling from the mother-bud remains suppressed till the nuclear migration is completed. After completing the migration and prior to the anaphase the metaphase spindle axis must become parallel with the mother-bud axis [8, 13, 54, 55, 60–62]. In Fig. 5f, we vary the position of septin ring in the hyphal tube and measure the nuclear migration time and the spindle orientation. We observe that finite migration time and parallel (almost) spindle orientation for all positions of the septin ring in the hyphal tube.

## DISCUSSION

The purpose of mitosis is to faithfully segregate the sister chromatids between the mother and daughter cells that are genetically identical [6, 56, 67, 79]. Mitosis consists of several phases and each of these phases must complete within a definite time frame [8, 94]. Earlier we identified mechanistic pathways and kinetics of various self-assembled chromosome-segregating machineries essential for pre-mitotic (interphase) and mitotic phases in budding yeasts [54, 55]. In the present study, we consider the yeast-hyphae morphology (e.g. in *S. Cerevisiae* and *C. Albicans*) to discern the mechanistic pathways functioning for the proper nuclear migration which is essential for mitosis. The destination of the nucleus is the septin ring that can localize anywhere inside the hyphal tube and since the hyphal tube can be greatly extended, the journey of the nucleus from the mother-bud is long [3, 25, 26, 29]. We quantify the mechanical forces arising from the interactions of dynamic microtubules with various intracellular objects that could be responsible for nuclear movement. These forces (viz., cortical pushing, pulling and sweeping) are transmitted to the nucleus through the astral microtubules [54, 55, 69]. Among these forces, we found that some promote migration (Fig. 2a and Fig. 3b) while the others suppress (Fig. 2c and Fig. 3a). When the migration-promoting forces are increased or suppressing forces are decreased, migration time is shorter. Since migration positions the nucleus at the site of division, it must conclude in a timely manner to facilitate proper mitosis. Delayed mitosis is error-prone that leads to improper segregation of the chromosomes and cell death [29, 94]. Our data show that irrespective of the hyphal length and the septin position in the hyphal tube, the nuclear migration is completed within a reasonable time [34] when dynein pulling from the mother-bud is insignificant (Fig. 5f). These results indicate the robustness of the mechanistic pathway as established by our *in silico* model.

A key difference between nuclear migration in yeast and hyphae is that in the first a shorter path is traversed by the nucleus, while in the latter the path is much longer through the hyphal tube [3, 5]. The nuclear movement in yeast remains restricted to the mother-bud as the septin ring is located at the junction between mother- and daughter-buds, whereas in hyphae, the nucleus travels through the germ tube (daughter-bud) [3, 5]. During yeast mitosis through the budding process, spatiotemporal characteristics of mechanical interactions among various molecular players are likely to differ between mother- and daughter-buds [8, 54, 68, 93]. For hyphae, the pattern of the interactions is unclear. The analysis from this study suggests a significant contribution of the cortical forces on migration. More specifically, a strong pulling force from the hyphal cortex must act on the nucleus. This is achieved by the astral microtubules from one of the SPBs sliding frequently along the hyphal cortex and engaging with the cortical dyneins. Irrespective of a fixed density or a stochastic variation, the interaction of dyneins with the cortical part of the microtubule generates a pull adequate for timely nuclear migration. In addition, our data shows the alignment of the mitotic spindle with the cell axis, indicating the cortical pathway’s robustness for successful mitosis.

A natural extension of the present model framework would be to investigate other possible scenarios for nuclear migration that explicitly focus on *“microtubule gliding”, “dynein ‘pull’ on nucleus”* and *“transport of nucleus as cargo”*. Recent studies suggest that dynein might act like a catch-bond while pulling on the microtubules [95, 96]. A catch-bond would allow dyneins to work at larger stall forces. Therefore, it would be worth exploring the nuclear migration dynamics considering stochastic dyneins in the light of catch-bond kinetics. Besides the timely migration of the nucleus to the site of division, another crucial factor determining mitosis’s fidelity is the correct attachment of the chromosomes to the spindle. An interesting future goal would be to examine the chromosome-spindle attachment where monotelic, syntelic and merotelic attachments between the kMTs and kinetochores can arise and propose a suitable “correction mechanism”. Also the role of spitzenkörper to nucleate astral microtubules or to guide them toward the septin ring could be checked. Suitable experiments could be designed to test the outcome of our simulation predictions which indicate frequent microtubule sliding in the hyphal cortex and suppressed dynein activities in the mother cortex for facilitating nuclear migration.

## ACKNOWLEDGMENTS

SS sincerely acknowledges the Indian Association for the Cultivation of Science (IACS), Kolkata, India for providing the fellowship and computational facilities. RP thanks IACS for the financial support and computational facilities.

## Notes

### Competing Interest Statement

The authors have declared no competing interest.

